# Untargeted ^1^H NMR Metabolomics and Pathway Analysis Reveals Dysregulated Proteostasis in Cyclophilin D (CypD)-deficient Mice Tissues

**DOI:** 10.1101/2021.04.20.440581

**Authors:** YI Adegbite, OS Adegbite, Y Ouyang, R Sutton, DN Criddle, LY Lian

## Abstract

Cyclophilin D (CypD) is a nuclear-encoded mitochondrial protein. Although best known as a regulator of the mitochondrial permeability transition pore (MPTP), it is also implicated in the regulation of cellular bioenergetics and extramitochondrial activities. In addition, being a peptidyl prolyl *cis-trans* isomerase (PPIase), deletion of CypD is likely to affect protein stability in the mitochondria; however, there is little direct evidence of this. In this study, untargeted ^1^H NMR metabolomics, coupled with multivariate analysis, was used to describe simultaneous changes in the metabolic system of CypD-deficient mice liver, heart, and pancreas, with data from the serum to identify systematic changes. Metabolomics Pathway Analyses (MetPA) revealed commonly perturbed metabolites in the different mouse tissues lacking CypD, with significantly enriched pathways that are related to amino acid, glucose and purine metabolisms, and bioenergetics. Serum from CypD-deficient mice confirmed changes in tissue urea cycle, lipid metabolism and ketogenesis. Overall this study reveals the role of CypD in maintaining protein homeostasis, concurring with its biochemical property as a peptidyl-prolyl isomerase and also demonstrates the wider metabolic adaptations induced by the deletion of the *ppif* gene, resulting in a CypD-deficient mice metabolome that is different from the wild-type system.

## Introduction

Cyclophilin D (CypD) is nuclearly-encoded by the *ppif* gene, and found in the mitochondrial matrix protein. It belongs to the cyclophilin family of the peptidyl prolyl *cis trans* isomerases (PPIase) ^1,2^. CypD possibly regulates mitochondrial function through helping with the folding of numerous target proteins, although very few reports have in fact demonstrated this functional aspect of CypD ^3^.

CypD is best recognised as a regulatory component of the mitochondrial permeability transition (MPT) pore, which is a non-specific channel whose proper function is key to mitochondrial health^4–7^. Transient opening of the pore is important for the maintenance of mitochondrial homeostasis^8^. However, under conditions of extreme cellular stress, persistent pore opening triggers the massive and unselective transit of molecules larger than 1500Da^9^; this depolarises the inner mitochondrial membrane, depletes mitochondrial ATP and ultimately leads to cell death^10^. Being the only genetically verified component of the MPTP system^4^, many studies undertaken using CypD KO animal and cell lines have concentrated on determining the functional relationship between CypD and the pore^11,12^. However, it is now clear that CypD itself is a master regulator of metabolism^13,14^ and deletion of *ppif* gene is expected to have very significant and wide-range effects which may be beyond the regulation of the MPTP, and acting through other proteins^15,16^. Hence, for understanding both MPTP regulation and additional biological functions of CypD, it is important to characterise the global metabolic effects of chronic CypD deficiency. Most pertinent are recent evidence that connects CypD to extramitochondrial activities, which includes its control of cellular metabolism ^17,18^. Menazza *et al*^17^ performed proteomics to characterise changes in protein expressions in CypD KO heart and concluded that branched chain amino acid and pyruvate metabolisms, and the Krebs Cycle are altered in this tissue. Tavecchio *et al*^18^ showed metabolic changes due to deletion of the *ppif* gene; profiles of the metabolome of WT and KO mouse embryonic fibroblasts revealed impaired β-oxidation and enhanced glucose metabolism when CypD was delected.

Metabolomics is a powerful and prospective tool for simultaneously defining global and tissue metabolic alterations in response to a variety of changes, including genetic ablation of specific genes; the method can provide a vast understanding of the metabolic effects of such changes^19–21^. In this study, ^1^H NMR metabolomics was used to systematically study organ-specific metabolic alterations in CypD KO mouse. This approach allowed simultaneous investigations of the global metabolic response of mouse liver, heart, and pancreas to CypD deficiency, in parallel with an elucidation of the metabolic signatures observed in the serum. Given that CypD is a PPIase and plays a role in the protein folding in the mitochondria, a focus here is on changes in amino acid metabolism. Indeed, this study shows that amino acids levels and products of amino acids metabolism are commonly perturbed across all the tissues and serum samples in the absence of CypD, providing evidence that CypD plays a role in maintaining proteostasis in the mitochondria.

## Results

### Metabolic profiling of samples

^1^H NMR spectroscopy was used to acquire the metabolic phenotype of CypD WT and KO mice tissues and serum. Approximately 45 and 88 individual metabolites were assigned to signals obtained from, respectively, serum and tissues of CypD WT and KO mice, using the metabolite libraries from Chenomx (https://www.chenomx.com/), the Human Metabolome Database (https://hmdb.ca/), and Biological Magnetic Resonance Data Bank (https://bmrb.io/), as well as in-house compound library which was verified by 1D and 2D spectroscopy. All the tissues shared the same metabolites (with the exception of dimethylamine which was not detected in the heart). Representative 1D ^1^H NMR CPMG spectra for the different tissue samples showed strong signals arising from water soluble metabolites such as amino acids, glycolytic metabolites, TCA cycle intermediates, metabolites of the urea cycle, one carbon units, nucleotide sugars, hexosamines, redox and energy co-enzymes (Fig. 1A-C). Spectra from the serum samples additionally contained signals arising from non-specific lipid/protein moieties observed as broad peaks and designated as mobile lipids at 0.81-0.91, 1.21-1.32, and 3.21-3.23 ppm spectral regions (Fig. 1D).

**Figure 1.**
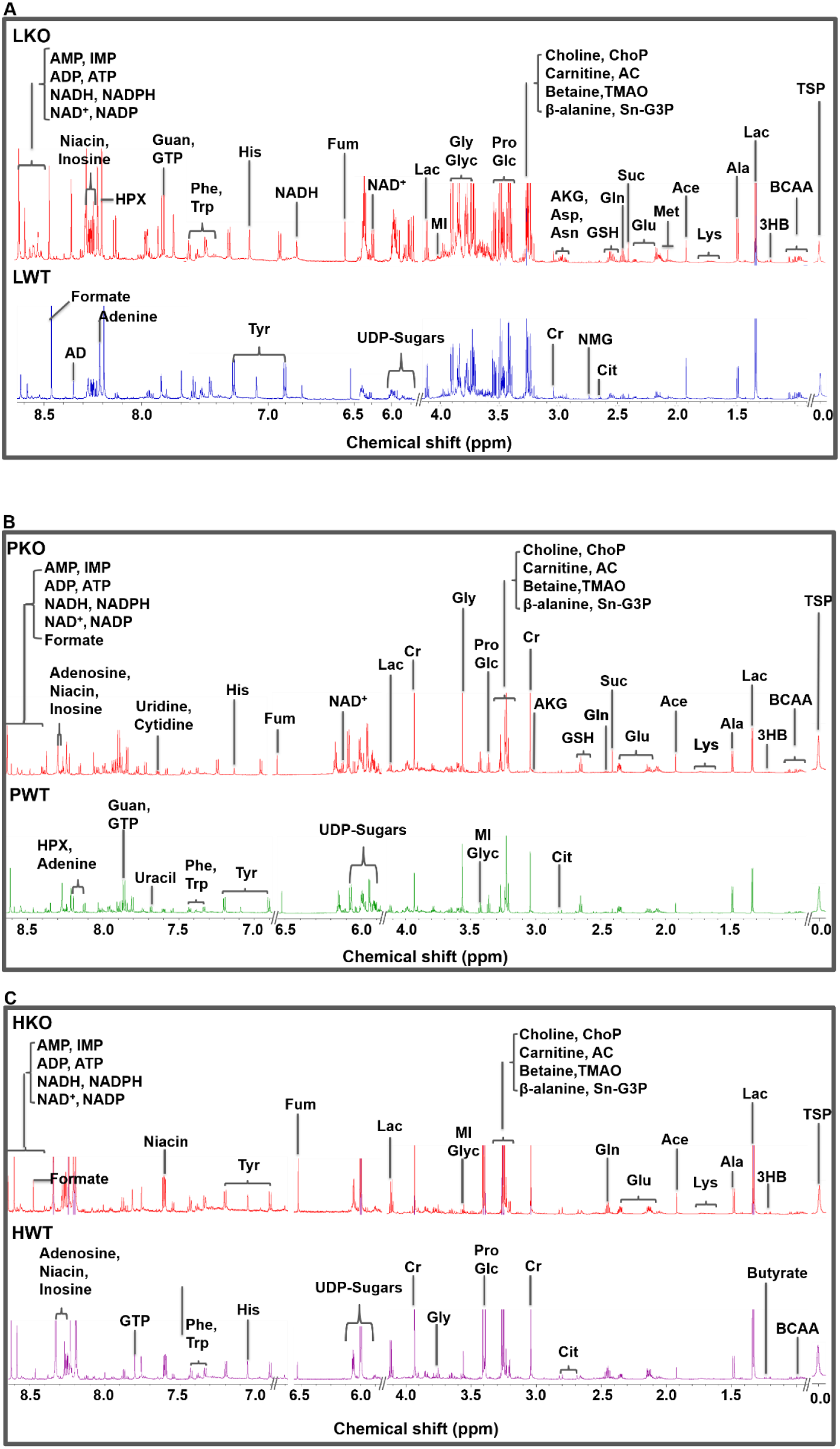
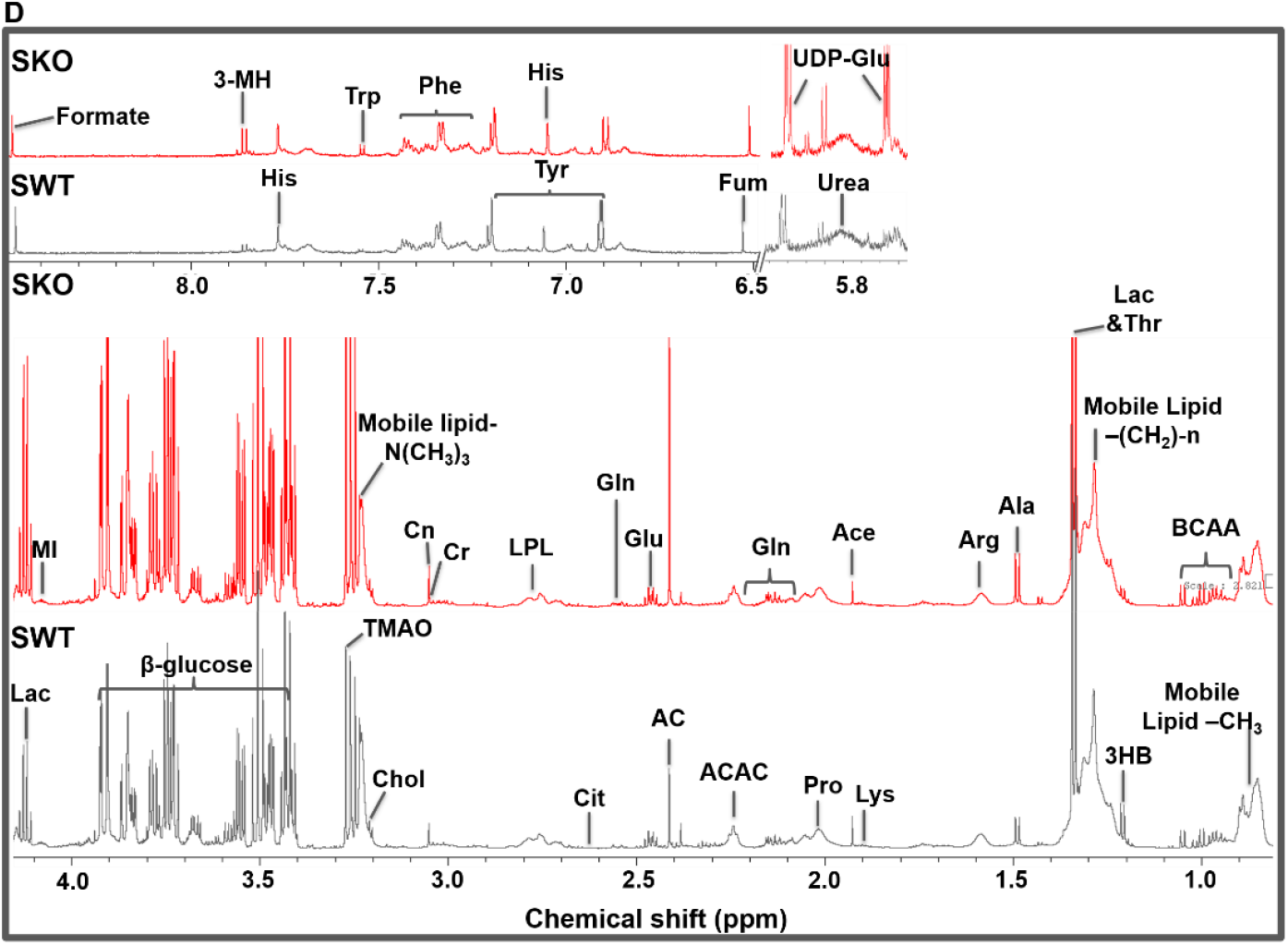
Representative 1D ^1^H NMR CPMG spectra of polar metabolic profile of CypD WT and KO extracts from **(A)** Liver **(B)** Pancreas **(C)** Heart and (D) Serum. All tissue spectra were referenced to 0.2mM TSP at 0.0 ppm. Abbreviations: BCAA, Branched Chain Amino Acids (Valine, Leucine & Isoleucine); 3HB, 3-hydroxybutryate; Lac, Lactate; Ala, Alanine; Lys, Lysine; Ace, Acetate; Glu, Glutamate; Suc, Succinate; Gln, Glutamine; GSH, glutathione; Met, Methionine, AKG, Alpha-ketoglutarate; Cr, creatine; PCr, phosphocreatine; PC, phosphocholine; sn-G3P, sn-Glycero-3-phosphocholine; Tau, Taurine; MeOH, Methanol; Glc, Glucose; UDP-sugars (UDP-glucose, UDP-glucuronate, UDP-galactose); Glc-6-P, Glucose-6-phosphate; Gal-1-P, Galactose-1-phosphate; Niacin, Niacinamide; Fum, Fumarate; His, Histidine; Tyr, Tyrosine; Phe, Phenylalanine; HPX, Hypoxanthine; AD, Adenosine; IMP, Inosine monophosphate; GTP, Guanosine triphosphate; AMP, Adenosine monophosphate; ATP, Adenosine triphosphate; ADP, Adenosine diphosphate, NADH, nicotinamide adenine dinucleotide (NAD)+ hydrogen (H); NAD, nicotinamide adenine dinucleotide; LKO, Liver KO; LWT, Liver WT; PKO, Pancreas KO; PWT, Pancreas WT; HKO, Heart KO; HWT, Heart WT; SKO, Serum KO; SWT, Serum WT.

All ^1^H NMR spectra were assessed by multivariate analyses. The first two components from unsupervised PCA, revealed a tight cohort clustering, indicating the reproducibility of the tissues and serum data, with liver demonstrating the highest impact of CypD deficiency on its metabolome when compared to the heart, pancreas, and serum (Fig. 2A-D). Independent Sample t-Test was used to screen mice tissues and serum metabolites that significantly discriminated between CypD WT and KO cohorts. Based on the false discovery rate (FDR) approach ≤ 0.05 and fold change ≥ 1.50 as cut-off, the level of 69, 55, 57, and 18 metabolites were significantly altered, representing 78.4%, 62.5%, 64.8%, and 40% of total measurable metabolites in, respectively, KO mice liver, heart, pancreas, and serum (Supplementary Figure 1). The list of metabolites with their corresponding chemical shifts, fold change and adjusted p-values for all tissue extract and serum samples of CypD WT and KO mice are provided in Supplementary Table 1.

**Figure 2:**
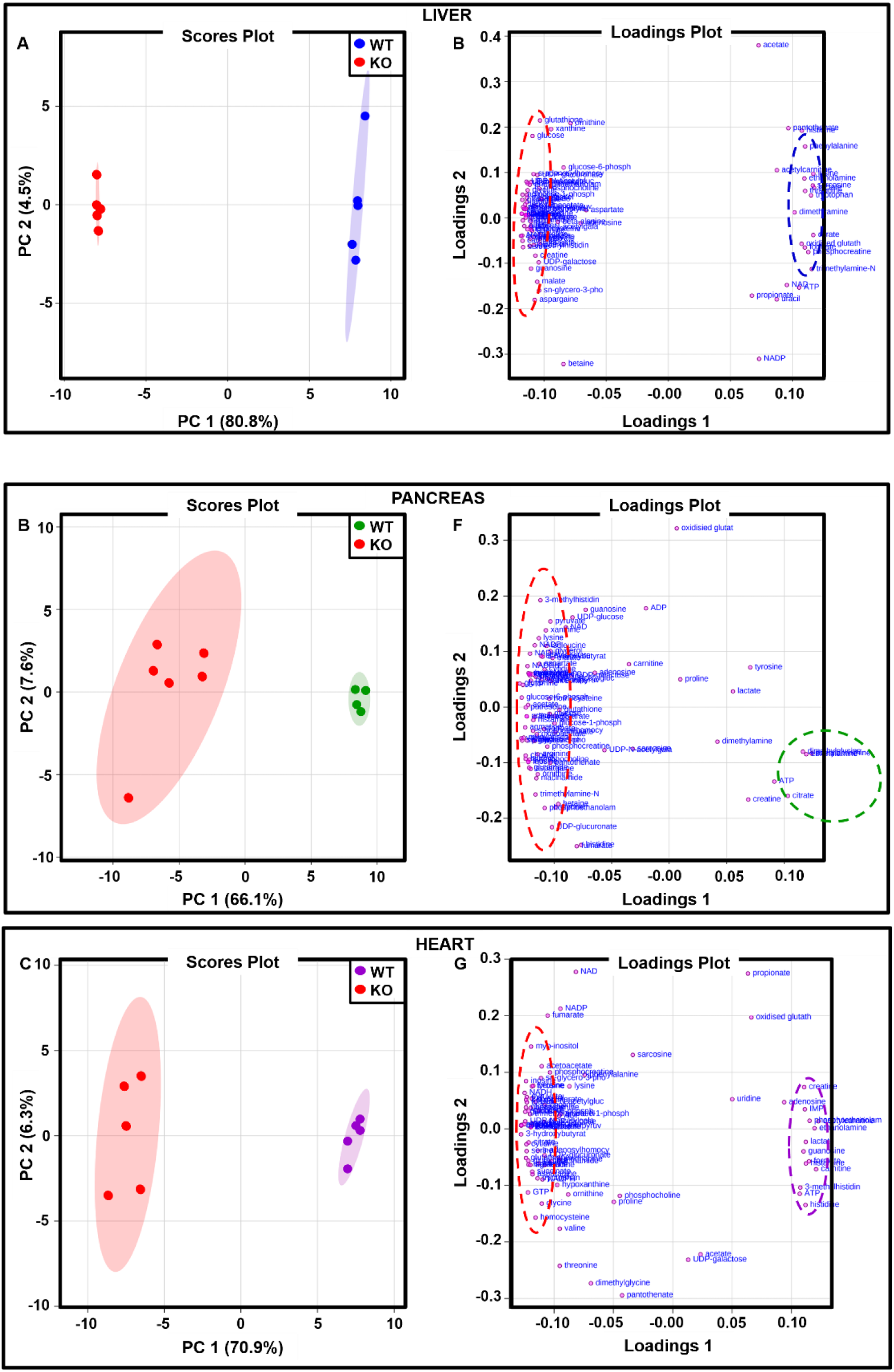
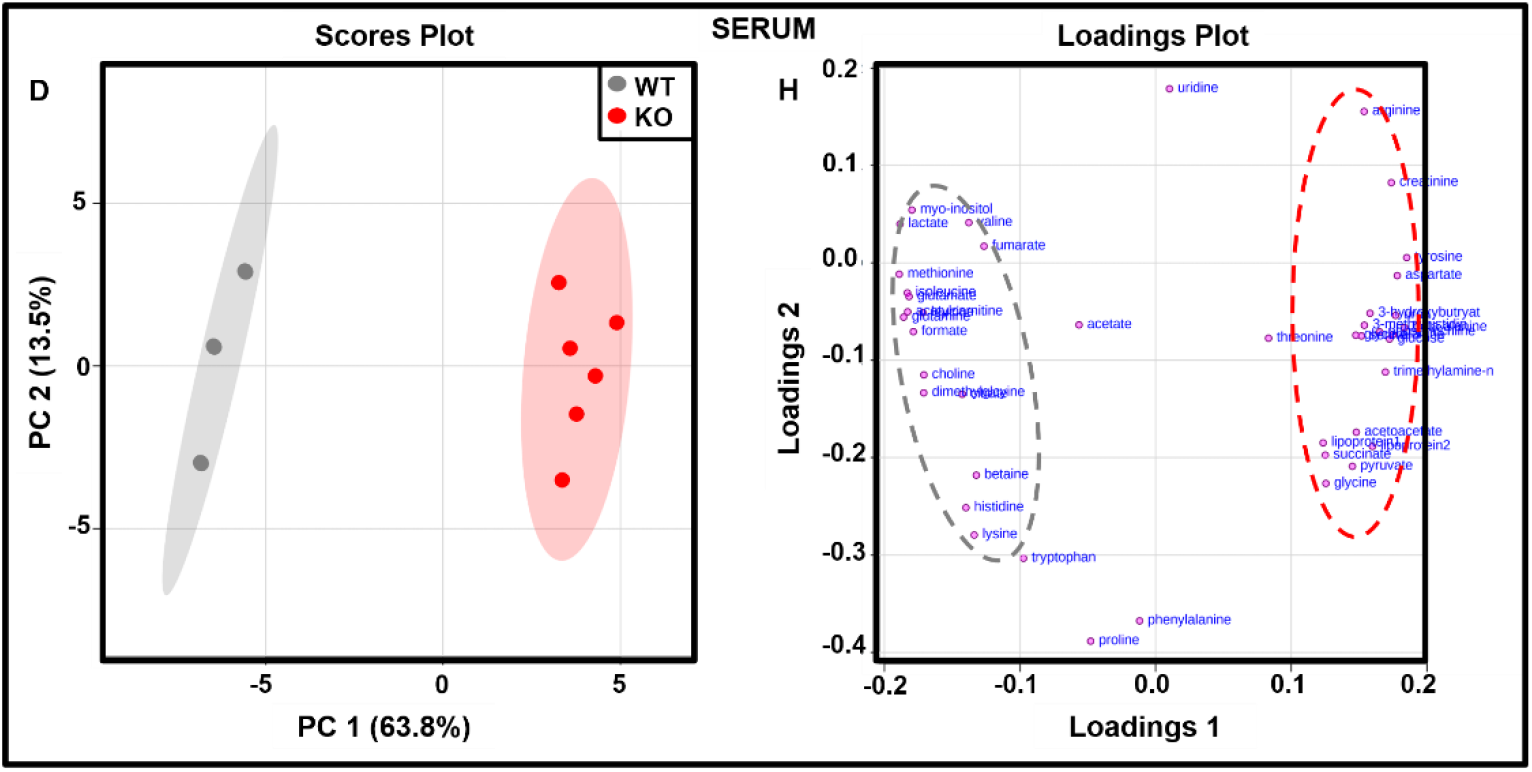
Two-dimensional scores plot **[A–D]** and loadings plot **[E-H]** from PCA of ^1^H NMR spectra for CypD WT and KO mice Liver **(A,E)**, Pancreas **(B,F)**, Heart **(C,G)**, and Serum **(D,H)**. Values in the axes refers to the total explained variance for each principal component represented in the scores plot. Ellipses represents 95% confidence. Most influential variables separating groups along the PC 1 axis only are highlighted in dashed ellipses for each mice tissues and serum in the loadings plots.

### Metabolite meta-analyses of CypD-deficient mice tissues and serum

To identify metabolites that were *commonly* impacted across the tissues and serum upon the genetic loss of CypD, meta-analysis was performed through the integration of individual datasets from mice tissues and serum. According to Fisher’s combined probability test, at a p-value significance threshold of 5%, a total of 28 metabolites were shared between the tissues and serum, a majority of which were amino acids, TCA cycle intermediates, and glucose and other glycolytic metabolites. Bar plots displaying the expression patterns of these commonly perturbed metabolites are shown in Fig. 3.

**Figure 3.**
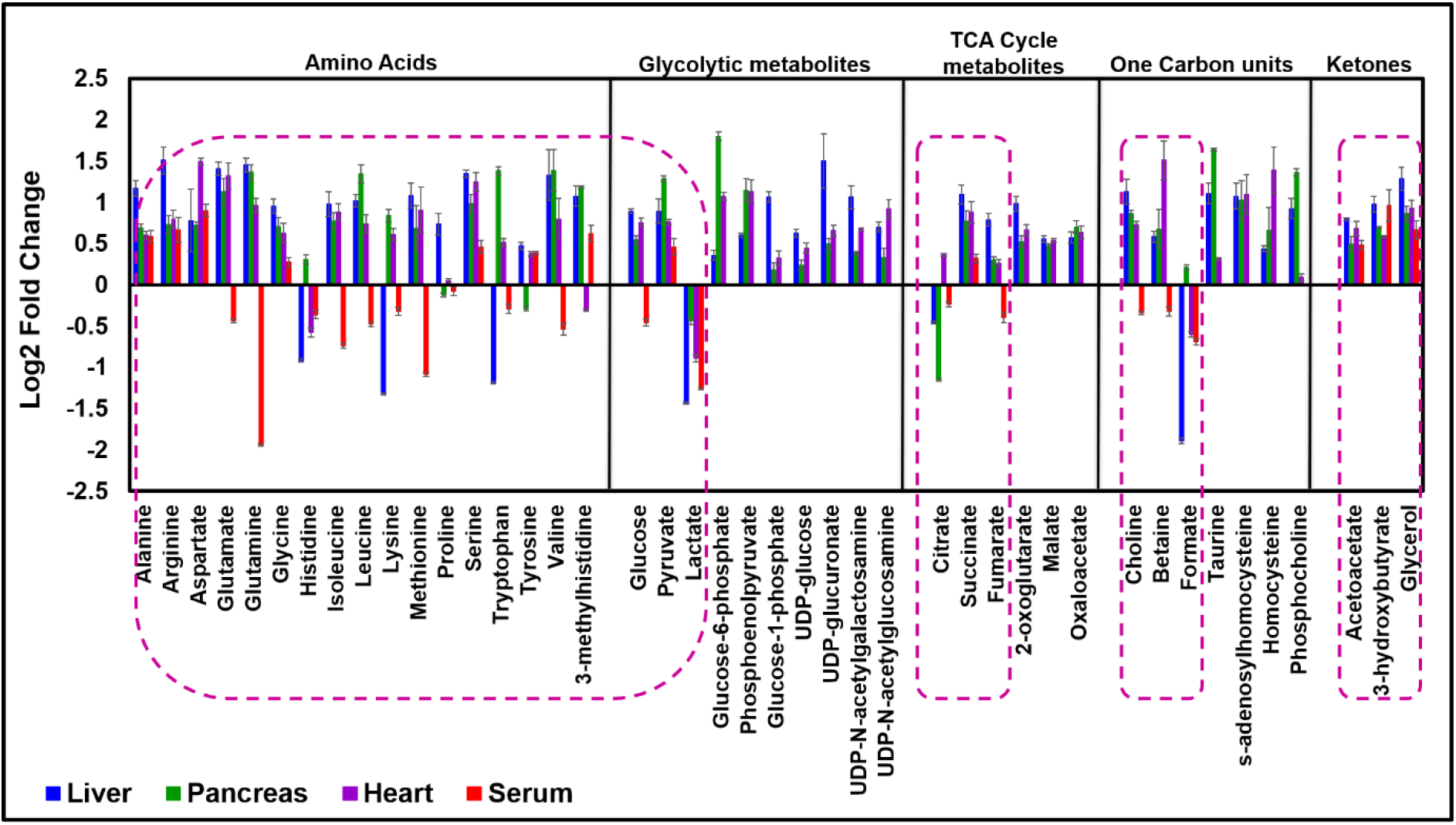
Bar plot displaying the Log2-fold change differences in the concentrations of commonly altered metabolites across CypD deficient mice tissues and serum with respect to the WT cohort. Bars facing the positive and negative log2 FC values respectively, depicts the increased and decreased metabolites in the KO tissues and serum relative to the WT. Only metabolites highlighted in dark pink dotted circles were universally altered across mice tissues and serum. Error bars depict standard error of the normalised mean for each compound. Statistical significance for each metabolite represented in this figure is *p* ≤ 0.05.

#### Amino Acids

Many amino acid levels were altered by CypD deficiency compared with Wt samples. Consistent across all mice tissues and serum was the elevation of some glucogenic amino acid (alanine, arginine, aspartate, glycine, and serine) while other glucogenic amino acids (methionine, glutamate, glutamine) and the branched chain amino acids (isoleucine, leucine, and valine) were also increased in all the KO tissues, but reduced in KO serum. The expression patterns of histidine, and the three ketogenic amino acids tyrosine, tryptophan, and lysine were altered in diverse manners with histidine increased KO pancreas but reduced in CypD deficient mice liver, heart and serum whereas the opposite was observed for tyrosine levels, being increased in KO mice liver, heart, and serum, but reduced in KO pancreas. Moreover, lysine and tryptophan concentrations were reduced in KO liver and serum, and elevated in mice heart and pancreas (Fig 3). Altogether, these observations illustrate that CypD deficiency affects amino acid levels on mice tissues and serum, albeit in a diverse manner. Of note was also an elevation of 3-methylhistidine in all but heart samples; this modified amino acid is a good marker for muscle protein degradation.

#### Metabolites of Glucose metabolism

The concentration of metabolites involved in glycolysis were also commonly perturbed in response to the loss of CypD in mice tissues and serum. Although mice tissues lacking CypD were characterised with lower levels of lactate, its upstream metabolites such as glucose, glucose-6-phosphate, phosphoenolpyruvate, and pyruvate were significantly increased. Likewise, the KO serum exhibited lower and higher concentrations, respectively, of lactate and pyruvate although KO serum glucose was lower (Fig. 3). In addition, increase in the levels of metabolites involved in glucose metabolism such as glucose-1-phosphate and UDP-glucose (involved in glycogenesis) were observed in the liver, but not significantly altered in other KO tissues. Finally, the concentrations of UDP-glucuronate (involved in glucuronidation), UDP-N-acetylglucosamine and UDP-N-acetylgalactosamine (part of the hexosamine biosynthetic pathway) were elevated in all the KO tissues when compared to their WT cohorts (Supplementary Table 1).

#### Metabolites of Tricarboxylic Acid (TCA) Cycle

Metabolites 2-oxoglutarate, succinate, fumarate, malate, and oxaloacetate were universally increased in all the KO tissues when compared to their WT cohort. Notably, citrate level was only higher KO heart tissues whereas both CypD-deficient mice liver and pancreas were characterised by reduced citrate concentrations. Though TCA cycle metabolites detected in the KO serum did not exhibit significant changes there was a modest decrease in citrate and fumarate levels by 1.31- and 1.17-fold, respectively, while succinate was increased by 1.24-fold in the KO serum (Fig. 3).

#### Metabolites of fatty acid metabolism

Metabolites associated with fatty acid metabolism such as 3-hydroxybutyrate and acetoacetate (ketones), and glycerol were increased in the KO tissues and serum when compared to their corresponding WT cohort (Fig 3).

#### Choline metabolites and One Carbon units

Metabolites involved in choline and methionine metabolism and in the transsulfuration pathway were consistently elevated in KO tissues. These metabolites are interlinked in the one carbon metabolic pathway for the *denovo* synthesis of amino acids, polyamines, purines, and phosphocholines. The level of choline, betaine, serine, glycine, methionine, formate, taurine, s-adenosylhomocysteine, and homocysteine were all significantly elevated in the KO tissues (Fig. 3). In addition, dimethylglycine which is derived from choline, is also high in KO liver and heart. All were consistently low in serum. Moreover, *sn*-glycero-3-phosphocholine (a precursor to choline biosynthesis and intermediate in phosphatidylcholine metabolism), was also significantly increased in the KO tissues (Supplementary Table 1). Choline and betaine, metabolites derived from diet, were reduced in the KO serum (Fig 3), while other metabolites of one carbon metabolism were not detected in the serum samples for analysis.

#### Independent metabolite analyses of CypD-deficient mice Tissues

Independent analysis was performed on metabolites that were *not universally* perturbed in the KO tissues and serum mice samples. These metabolites include the purines, redox and energy associated metabolites, which were not detected in the serum samples. Box plots displaying the abundance differences between the WT and KO cohorts are represented in Fig. 4.

**Figure 4.**
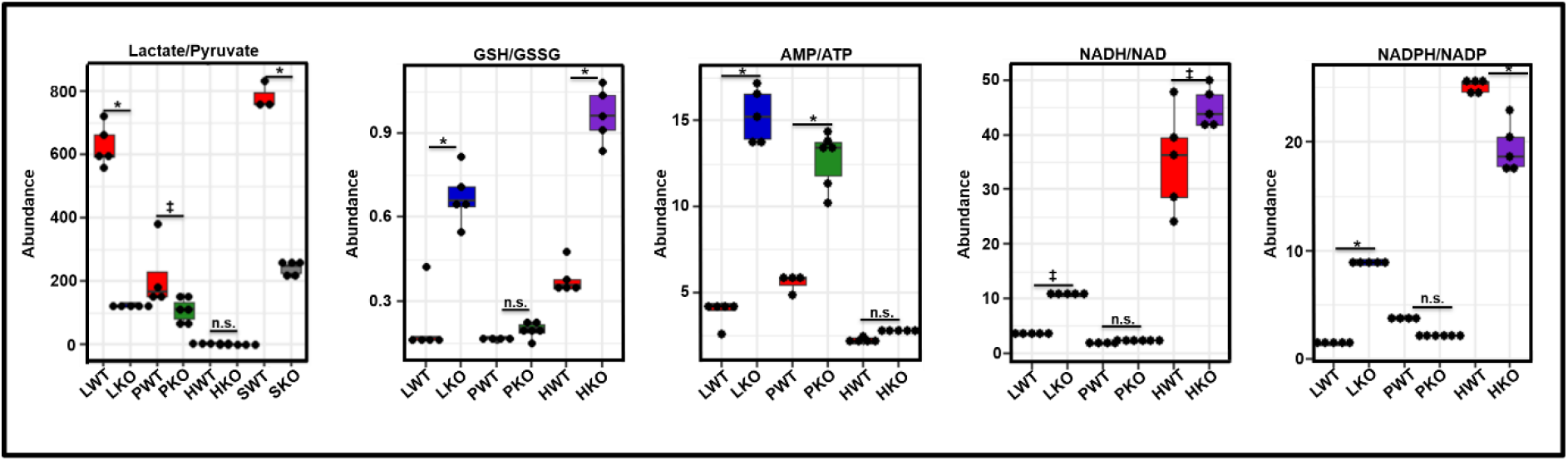
Metabolic flux of CypD rich and deficient mice tissues and serum. Metabolic flux was calculated by the ratio of two interconvertible metabolite concentration. The ratios suggest the possible metabolic flow between CypD WT and KO mice tissues and serum. Statistical significance was determined by ANOVA and post hoc Tukey’s HSD, using R programming. \**p* ≤ 0.0001; ^†^*p* ≤ 0.001; ^‡^*p* ≤ 0.05; n.s. means not statistically significant. Data represented as mean ± SEM. Abbreviations: LWT, Liver WT; LKO, Liver KO; PWT, Pancreas WT; PKO, Pancreas KO; HWT, Heart WT; HKO, Heart KO; SWT; Serum WT, SKO; Serum KO.

#### Redox co-enzymes and Bioenergetics

The absence of CypD in mice liver, heart, and pancreas perturbed the level of metabolic biomarkers for redox defense e.g. the concentration of NADH and NADPH were significantly elevated in all the KO tissues relatively to the WT cohort. In the liver, this effect was accompanied by a significant drop in the level of NAD^+^ and NADP^+^ in this tissue, while the KO pancreas and heart demonstrated elevated concentration of these compounds (Supplementary Table 1). Increased ratios of NADH/NAD^+^ in the CypD deficient mice liver and heart were also observed. The ratio of NADPH/NADP^+^ was elevated in the KO liver and decreased in the KO heart. Moreover, the concentration of glutathione (GSH) was significantly higher in the KO tissues while its oxidised form (GSSG) was reduced in the KO liver and pancreas, but unchanged in the KO heart, leading to observed significant increases in the GSH/GSSG ratio in the KO liver and heart (Fig. 4); this suggests that overall, CypD-deficient mice tissues may be characterised with a reductive environment.

The genetic loss of CypD also altered the bioenergetics of mice tissues, as evidenced by the decreased and increased levels of, respectively, ATP and AMP in the KO, with respect to their corresponding WT cohorts, leading to an increase in the AMP/ATP ratio, and indicating a fall in the energy status of the KO cells (Fig. 4).

#### Purines

The concentrations of purines such as hypoxanthine, xanthine, inosine, adenine, and GTP were found to be elevatedr in the KO tissues. The expression patterns of other detected purines such as IMP and ADP were, however, altered in a divergent manner. In comparison to their WT counterpart, the level of IMP was lower in the KO heart but increased in the KO liver and pancreas, whereas the level of ADP was higher in KO liver and heart but was not altered in KO pancreas (Supplementary Table 1).

#### Independent metabolite analyses of CypD-Deficient mice serum

Overall fewer metabolites were observed in the serum samples; notably missing were many of the redox and coenzyme metabolites (NAD^+^, NADH, NADP^+^, NADPH, GSH, and GSSG) as these are unstable and occur at low concentrations in the serum. Nevertheless, CypD-deficient mice serum showed changes in similar classes of metabolites as found in the tissue samples; in some cases, the metabolite changes were in the similar direction, but reverse in others (Fig. 3, Supplementary Table 1). Notably, serum-specific metabolites not detected in the tissues such as urea (a by-product of protein degradation and amino acid metabolism), creatinine (a by-product of muscle metabolism), circulating mobile lipids (representing low density (LDL), and very low-density lipoproteins (VLDL)), were significantly accumulated in the KO when compared to the WT cohort (Fig. 5).

**Figure 5.**
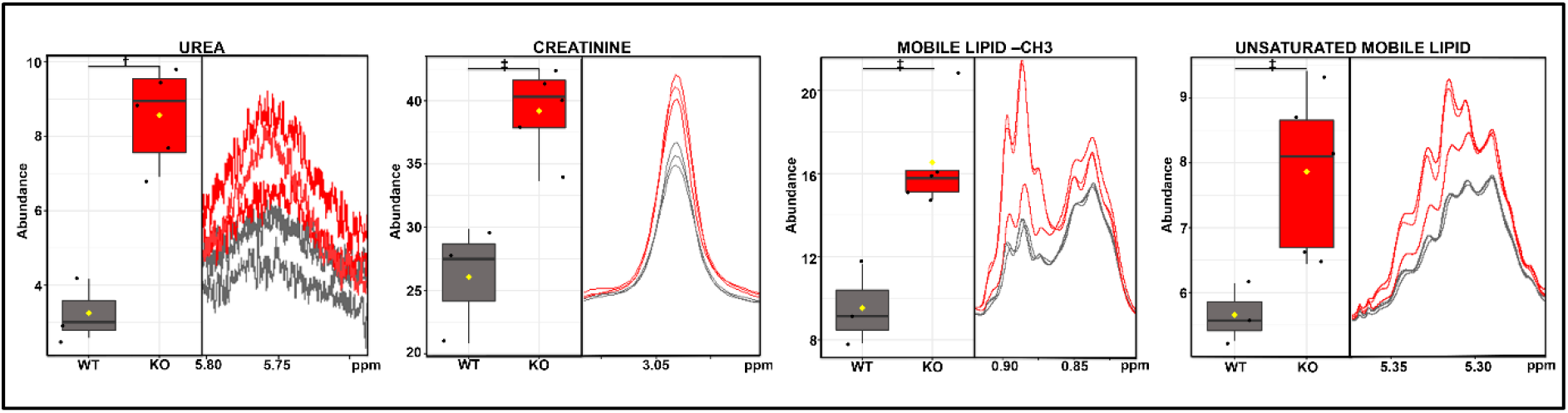
The abundance and distribution of serum specific metabolites discriminating between CypD WT and KO mice serum. Boxes representing each metabolite shows the box plot (left panel) and its corresponding overlayed (3 samples per cohort) 1D ^1^H NMR CMPG spectra (right panel). The black dots represent the normalised concentrations of the metabolite in each cohort. The horizontal black line denotes the median value while the mean concentration of each cohort is indicated with a yellow diamond. Statistical significance for each metabolite is represented with a statistical sign: \**p* ≤ 0.0001; ^†^*p* ≤ 0.001; ^‡^*p* ≤ 0.05. n = 3 (WT) and n = 5 (KO).

### Metabolic pathway analysis of CypD WT and KO mice tissues and serum

Pathway analyses using the “Metabolomics Pathway Analysis” (MetPA) software package from Metaboanalyst 4.0 were performed on metabolites that were consistently altered between CypD WT and KO cohorts, at a statistical significance value ≤ 0.05. In CypD-deficient mice tissues, significant alterations in 30 metabolic pathways, of which 26 were impacted at a score ≥ 0.1 were deduced; the other 4 pathways with an impact score < 0.1 were considered not biologically affected in response to the loss of CypD.

Pathways that were significantly affected, with an impact score ≥ 0.1, by the loss of CypD were mainly amino acid-related pathways (e.g. valine, leucine and isoleucine degradation, alanine, aspartate, histidine, arginine+proline, glutamate, phenylalanine+tyrosine, glycine+serine, and methionine metabolism) and mitochondrial associated pathways (e.g. transfer of acetyl groups into the mitochondria, malate-aspartate shuttle, betaine metabolism, TCA cycle, pyruvate metabolism, Warburg effect, mitochondrial electron transport chain, folate metabolism, glycerolipid metabolism, urea cycle and ammonia recycling). Other metabolic pathways that were also significantly impacted in response to CypD-deficiency are metabolic pathways related to glucose metabolism e.g. glucose-alanine cycle, glycolysis, and gluconeogenesis. Finally, purine metabolism (enriched by amino acids, nucleotides, and nucleosides) was also significantly affected as a result of CypD deficiency, with an impact score of 0.18.

In the case of the serum metabolic profile, only beta-alanine metabolism, ammonia recycling and urea cycle were able to reach statistical significance. Figures 6A and B are the graphical outputs of the significantly altered metabolic pathways which were generated from the Small Molecule Pathway Database (SMPDB), based on “normal human metabolic pathways”; a more detailed list is provided in Supplementary Table 2.

**Figure 6.**
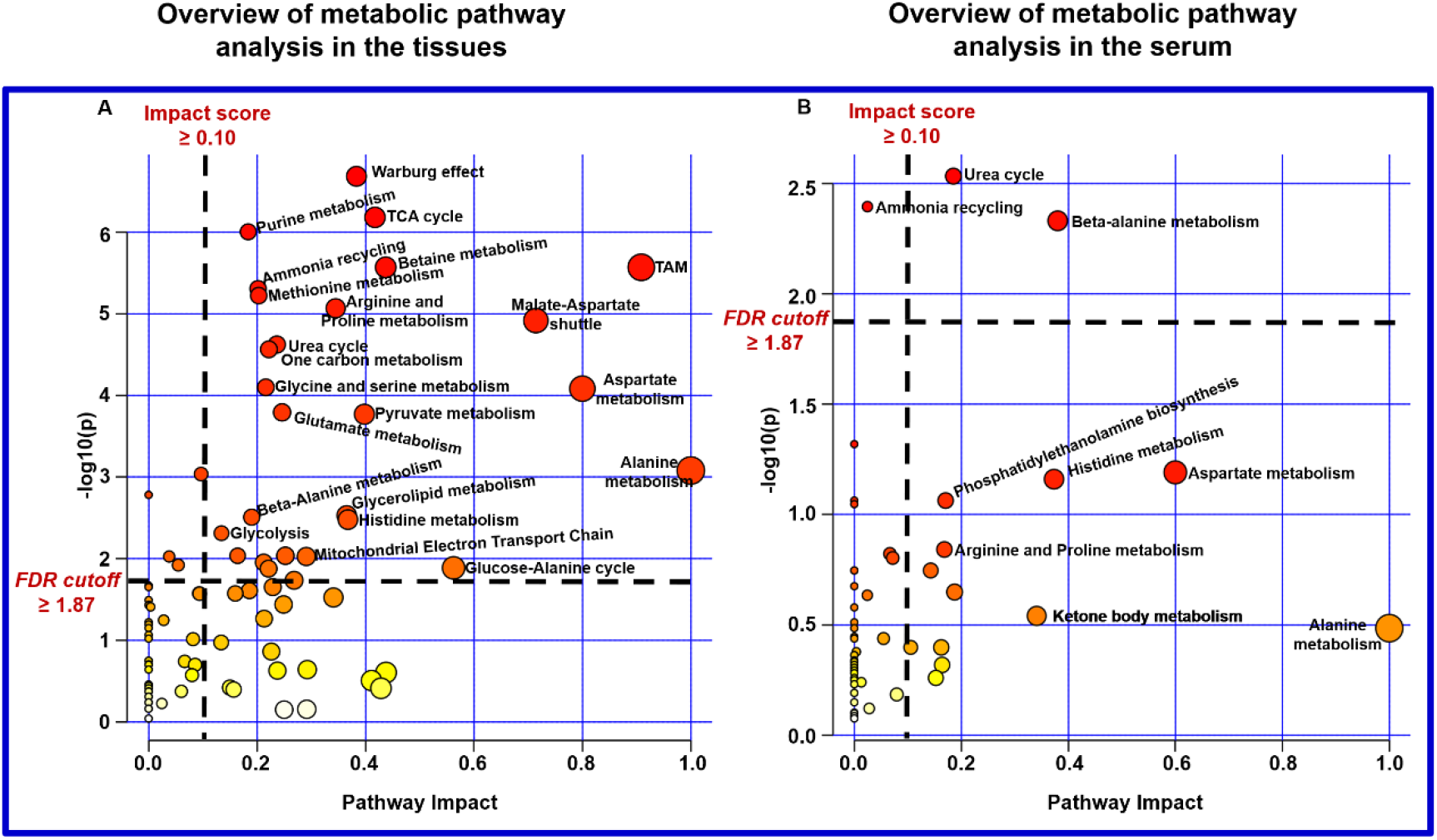
Graphical visualization of the metabolome view of the pathway enrichment and topology analysis of significantly altered metabolites from CypD WT and KO mice **(A)** tissues and **(B)** serum. Pathway Score in x-axis represents the impact of these enriched pathways computed from topology analysis, while −log10(P) in y-axis refers to the negative natural logarithmic value of the original p-value from statistical analysis of pathway difference between the two cohorts. Data points above the vertical black dotted line are statistically significant (–Log false discovery rate ≥ 1.87, is equal to p ≤ 0.05). Data points below the horizontal black dotted line to the right side of the graph, are acceptably impacted (Impact score ≥ 0.1). Each circle represents a different pathway; circle size and color shade are based on the pathway impact and p-value (red being the most significant), respectively. Abbreviations: TAM, Transfer of Acetyl cohorts into the Mitochondria.

## Discussion

The objective of this study was to unravel the universal role of CypD in the control of cellular metabolism by performing ^1^H NMR metabolomics on CypD KO tissues and serum samples. The results provide a holistic view of the effects of CypD deficiency since the different tissues explored possess different basal metabolic needs and physiological functions. Interestingly, amongst the three tissues studied, CypD is most abundant in the heart, and lowest in the liver (unpublished data); yet the liver showed the largest changes in metabolite characteristics. Explanations for some of the observed metabolite changes could be found in the widespread changes in protein expression in heart tissues resulting from CypD KO which were reported by Menazza et al^17^.

Overall, the deletion of CypD altered a wide range of metabolites in mice liver, heart, and pancreas. For some metabolites, the tissues showed opposite effects from each other upon CypD deletion whereas other metabolites were found to be altered only in specific tissues. An important example of opposite effects is the NADPH/NADP^+^ ratio of liver, heart and pancreas in response to CypD deficiency. Compared to WT mice, KO liver displayed elevated NADPH/NADP^+^ ratio, thus increasing the naturally occurring biosynthetic potential of liver tissues, whereas this ratio was reduced in both pancreas and heart tissues.

Of note are two important metabolites, citrate and formate, which are both produced in the mitochondria; citrate from the TCA cycle, and formate, through catabolism of serine and sarcosine. The levels of both were reduced in KO liver, hence, directly reflecting changes in mitochondria metabolism in hepatocytes. On the other hand, the rise in citrate and fall in formate levels observed in CypD KO heart, and the reverse in CypD KO pancreas, were likely the result of differences in activities and abundances of the associate metabolic enzymes. A schematic representation of the metabolites and metabolic pathway alterations in CypD-deficient mice serum, liver, heart, and pancreas is shown in Fig. 7.

**Figure 7.**
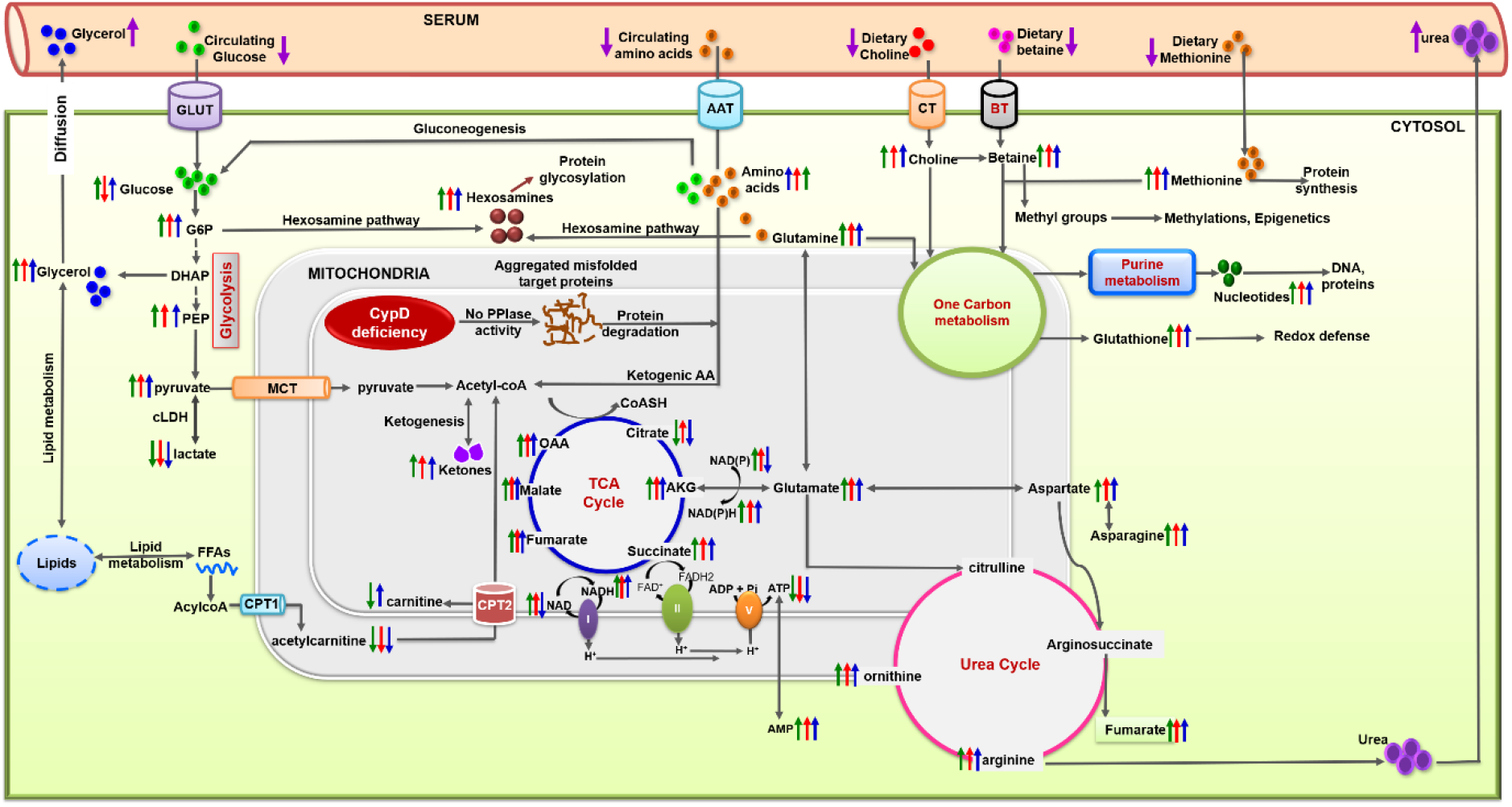
Schematic representation of the metabolites and metabolic pathway alterations in CypD-deficient mice serum and tissues: liver, heart, and pancreas. Only metabolites and pathway of biological relevance to the genetic loss of CypD is illustrated in this figure. Arrows pointing in the upward and downward direction, respectively indicates significantly increased and decreased metabolites in CypD-deficient mice serum and tissues, relative to their WT cohort. Blue, Red, Green, and Purple arrows represent changes observed in, respectively, CypD-deficient mice liver, heart, pancreas, and serum. Abbreviations: GLUT, Glucose transporter; AAT, Amino acid transporter; CT, Choline transporter; BT, Betaine transporter; MCT, Monocarboxylate transporter; CPT1, Carnitine palmitoyltransferase 1; CPT2, Carnitine palmitoyltransferase 2; LDH, Cytosolic Lactate dehydrogenase; mLDH, Mitochondrial Lactate dehydrogenase; AKG, Alpha-ketoglutarate; OAA, Oxaloacetate; PCr, Phosphocreatine.

The metabolic profile of the KO serum showed distinct metabolome changes compared to KO tissues which is indicative of differences between systemic and localised responses to the deletion of the *ppif* gene (Supplementary Table 1). However, since blood is an important tissue of the circulatory system that allows for the efficient exchange of nutrients between organs^22^, it is reasonable to assume that metabolic changes found in the serum reflect metabolic changes taking place in mice tissues, in particular the liver, as shown in Figure 3.

The significant elevation of ketones (3-hydroxybutyrate and acetoacetate) and urea cycle metabolites (such as arginine, ornithine, urea) observed in the KO tissues was also found to be increased in the KO serum. Of the three solid tissues examined, KO liver displayed the highest fold changes in these compounds (Fig. 3); this is not surprising as the liver is the central hub of metabolism and biosynthesis of metabolites conveyed to other tissues via the serum^22^. On the other hand, unlike tissues, KO serum had reduced levels of many amino acids (e.g., glutamate, glutamine, isoleucine, leucine, lysine, methionine and valine, lysine, methionine) and other circulatory metabolites (e.g., betaine, choline, and myo-inositol), following the loss of CypD (Fig 3, Supplementary Table 1). This may be largely attributed to an overall increase in net influx/efflux ratio of amino acids in mammalian tissues^23^. Because amino acids are essential for metabolism in all tissue types and their tissue abundance is dependent on either absorption from the circulatory system or *in situ* amino acid metabolism/proteolysis, their reduced levels in the serum relative to tissues is to be expected.

Altogether, these data show that tissue-specific alterations in metabolites levels as a result of CypD deficiency are a consequence of differences in the physiological functional of these organs. These differences are determined by macromolecular contents such as proteins, carbohydrates, lipids and nucleic acids specifically located in the different organs^24^.

### CypD regulates amino acid metabolism

Several amino acid-related pathways are identified as universally impacted and enriched by CypD deficiency in mice liver, heart, and pancreas (Supplementary Table 2). They concur with previous proteomics studies of CypD deficiency by Menazza et.al ^17^, where a large number of proteins involved in branched chain amino acid degradation were found to be altered in CypD-deficient mice heart. Tissue amino acid homeostasis is regulated by the net effect of protein and amino acid synthesis or degradation, coupled with activities of amino acid transporters^24^. Elevated amounts of amino acid in the CypD-depleted tissues were consistently observed here. It is likely that this is due to mitochondrial stress associated with the loss of CypD peptidyl prolyl isomerase activity^13, 15^ resulting in dysregulated folding of client proteins. Indeed, relative to WT tissues, markers of the urea cycle, such as arginine and ornithine, were significantly upregulated in the KO liver, as well as in the heart and pancreas tissues where urea cycle pathway are not abundant under basal conditions (Fig. 4, Supplementary Table 1). Another marker of protein degradation, 3-methylhistidine, was also elevated all tissues except the heart.

In the KO serum, the reduced levels of the essential amino acids and metabolites commonly associated with diets such as betaine and choline (Fig. 3, Supplementary Table 2) may likewise reflect the need for individual tissues to replenish damaged mitochondrial proteins, this requiring energy. The aggressive uptake of available serum amino acids and other metabolites by the tissues is a compensatory mechanism for energy production and protein synthesis. The elevated urea level observed in KO serum emphasized enhanced proteolytic activities and protein turnover in these tissues, and this is likely to predominantly occur in the liver where basal urea cycle is high.

### CypD deficiency enhances glucose metabolism in mice tissues

The tissue of most interest here is the liver due to its central role in glucose metabolism. Excessive glucose and its intermediates such as glucose-6-phosphate, UDP-glucose, phosphoenolpyruvate and pyruvate present in the KO liver tissues suggest increased activities in one or more of the glucose pathways of glycolysis, gluconeogenesis and, possibly, glycogenesis. Increased glycolysis is supported by the elevated reductive equivalents present in KO tissues such as NADPH/NADP^+^ (liver tissues only) and NADH/NAD^+^ (other tissues). In addition, decreased intracellular lactate concentration and lactate/pyruvate ratio in all KO tissues including serum (Fig. 3,4, and Supplementary Table 1) suggest a reduced activity of lactate dehydrogenase, possibly in preference of gluconeogenesis to glycolysis pathways. Excessive serum and hepatic glycerol, a glucose precursor, was observed in KO mice. Since most extrahepatic tissues lack gluconeogenic capabilities, glucose obtained from gluconeogenesis in liver tissues is secreted into the serum to nourish other tissues. However, elevated hepatic glucose can also enrich the glycogenic store and other glucose metabolic pathways^25,26^. Indeed, glycogenic metabolites such as glucose-1-phosphate and UDP-glucose were found to be of higher concentration in CypD-deficient mice liver, when compared to the WT cohort whereas these compounds were not significantly altered in other tissue types (Supplementary Table 1). This is not surprising, because the liver is the main site for glycogenesis as well, while many other mammalian tissues lack this ability, as they are mainly consumers^27,28^. Finally, the elevated levels of glucogenic amino acids such as alanine, arginine, asparagine, glutamate, glutamine, glycine, methionine, proline (liver), serine, valine, phenylalanine (pancreas), isoleucine, threonine (pancreas, heart), tryptophan (pancreas, heart) and tyrosine (liver, heart), possibly arising from protein degradation, were also abundant in KO tissues, supporting the notion of increased glucose biosynthesis upon deletion of the *ppif* gene.

Interestingly, the intermediates of the hexosamine pathway, which is a minor branch of glycolysis (UDP-N-acetylglucosamine and UDP-N-acetylgalactosamine), were significantly higher in all KO solid tissues (Supplementary Table 1). Indeed, pathway analysis revealed that hexosamine metabolism were significantly impacted in the KO tissues.

### CypD deficiency perturbs one carbon metabolic pathway

Metabolic pathways that generate one carbon (1C) unit for the synthesis of macromolecules such as DNA and phospholipids, and other metabolites such as amino acids, polyamines, and purines were more abundant in all KO tissues than their WT. Interestingly, amino acids (e.g., serine, glycine, methionine, dimethylglycine, and betaine), polyamines (e.g., putrescine, histamine, and agmatine) and phospholipid components (e.g., phosphocholine, ethanolamine, and choline), were all elevated in the KO tissues than their corresponding WT groups. There was also an accumulation of S-adenosylhomocysteine, homocysteine, taurine, and glutathione, all of which are intermediary products of pathways linked to 1C metabolism (Fig 3, and Supplementary Table 1). The pathways affected include arginine + proline metabolism, and glycine +serine metabolism. The biological importance of the 1C metabolic reactions lies in its ability to support, through the donation of methyl groups, multiple physiological processes, maintain epigenetics and redox defense^29^.

Since reduced serum levels of choline and other amino acids alongside elevated intracellular concentrations in KO mice were observed, it is plausible that KO tissues are characterised by enhanced amino acid and phospholipid recycling, coupled with chromatin remodelling, which alters the overall signalling pathways of cells within these tissues.

### CypD regulates mitochondrial metabolism (TCA Cycle and Ketogenesis)

Elevation of 2-oxoglutarate, succinate, fumarate, malate, and oxaloacetate levels in all the KO solid tissues were observed (Fig. 3). However, there were differences in the citrate level changes for the KO tissues, this being reduced in the liver and pancreas but elevated in the heart. Since the synthesis of citrate from condensation reaction between oxaloacetate and acetyl-coA is the committed step of the mitochondrial TCA cycle, reduced citrate levels in the KO liver and pancreas suggest that for these tissues, unlike the heart, the initiation step of the TCA cycle was downregulated and concurred with the observed accumulation of pyruvate and alanine in these tissues

There were also elevated levels of acetoacetate and 3-hydroxybutryrate in all the KO tissues and serum (Fig 3). Ketone bodies are produced primarily in liver tissues and exported to be used in heart and other tissues, where they are converted to acetyl-CoA prior to TCA metabolism^30^. The elevated liver ketones in KO are most likely generated from fatty acids and amino acids, the latter supported by the increased levels of many ketogenic amino acids (see above). Increased ketone bodies levels in KO pancreas and heart tissues indicates the accumulation of acetyl-CoA in these extrahepatic tissues KO tissues. Unfortunately, acetyl CoA could not be definitively identified in the ^1^H NMR spectra of the samples due to signal overlap (Supplementary Table 1).

As far as the TCA cycle itself, the high energy demand observed in KO tissues results in enhanced alternative anaplerotic reactions of the cycle such as glutaminolysis, i.e. the glutamine/glutamate/2-oxoglutarate axis. Indeed, higher levels of 2-oxoglutarate and succinate were found in all KO solid tissues.

### CypD regulates Lipid metabolism

Glycerol, in this study, was used as the major marker for lipid metabolism^31,32^. Figure 3 shows increased levels of glycerol in KO serum and tissues. Glycerol homeostasis is regulated by lipid degradation and the glycolytic pathway^31,32^. Under conditions of high glucose availability, there is a steady supply of glycerol-3-phosphate from the reduction of dihydroxyacetone phosphate, a glycolytic intermediate, which often results in the synthesis of glycerol mainly in adipocytes and liver^31,33,34^. Adipose glycerol are secreted into the blood, and this is absorbed by the liver for glucose synthesis and triglyceride storage. Since glycerol and its derivatives are essential for lipid synthesis, it is plausible that lipid synthesis is elevated in the KO tissues, most especially lipogenic liver tissues. Interestingly, an elevation of mobile lipid signals in the KO serum (Fig. 4) was detected. It is likely that lipids synthesized in liver tissues and adipocytes were effectively exported into the serum for metabolism by other lipid metabolizing tissues such as heart tissues^34^. Interestingly, CypD deficiency has also been previously reported to impair fatty acid beta oxidation by lowering the expression level and activity of carnitine palmitoyl transferase I (CPT1), ultimately reducing acylcarnitine levels in mice heart^17^, and mouse embryonic fibroblasts^18^.The lower levels of intracellular and serum acetyl carnitine (the shortest form of acylcarnitine) detected here in CypD KO mice concur with these previous studies.

### CypD deficiency perturbs cellular bioenergetics and upregulates denovo nucleotide synthesis

Numerous amino acid substrates and one-carbon units such as glutamine, glycine, aspartate and formate, contribute to the *de novo* biosynthesis of purines^35–37^. Hence metabolic dependencies on glutamine and aspartate fuel the nucleotide synthesis pathway in KO tissues. CypD deficiency was characterised by elevated purine nucleotides such as xanthine, hypoxanthine, IMP, AMP, and GTP, and purine nucleosides such as inosine, adenosine, and guanosine (Supplementary Table 1). Since sufficient energy supply is required for the sustenance of some anabolic pathways present in CypD depleted tissues (especially hepatic tissues), it is plausible to assume that *denovo* nucleotide synthesis compensates for reduced mitochondrially generated ATP observed in the KO. Moreover, significant increase in other energy generating pathways such as cytosolic aerobic glycolysis (mainly heart and pancreas), mitochondrial glycerol-3-phosphate dehydrogenase-dependent respiration and glutamine anaplerotic TCA intermediate supply, further suggests the requirement to keep the cellular energy currency high to meet the metabolic demands in KO cells. Indeed, both NADH/NAD ratio (Fig. 4), and glycerol and its intermediates (*sn*-glycero-3-phosphocholine) were elevated in KO tissues relative to their WT counterpart (Supplementary Table 1). These observations suggest possible dysregulation of the mitochondrial respiratory chain in the absence of CypD, hence, the accumulation of reductive equivalents in our studies.

## Conclusion

^1^H NMR metabolomics studies on three key mammalian tissues - liver, heart and pancreas- and on serum were used here to unravel the effects, on the metabolome, of deleting the *ppif* gene which encodes the mitochondrial peptidyl-prolyl isomerase, CypD. The liver showed the most changes between CypD WT and KO samples, and serum the least. Most notable are the effects of CypD deletion on amino acid metabolism. The data supports the expected role of CypD in regulating mitochondrial protein homeostasis: deleting CypD resulted in universal changes in amino acid in three tissues and serum with significant impact on the urea cycle, indicating dysregulated proteolysis and protein turnover. The other significant pathways affected by loss of CypD relate to glucose and lipid metabolism, the TCA cycle and bioenergetics regulation; these imply perturbation of mitochondrial energy production and compensation from both TCA anaplerosis and *denovo* nucleotide synthesis, possibly due to deficient protein function and folding resulting from the absence of CypD. This perturbation of mitochondrial metabolism in turn elicits further adaptive response in which more widespread modulation of the expression of other metabolic enzymes lead to the eventual metabolome of CypD KO being different from the WT mice.

In summary, identifying the role of CypD in maintaining protein homeostasis concurs with its biochemical property as a peptidyl-prolyl isomerase, and provides an overarching explanation as to why the deletion of CypD, which exist is relatively low abundance in normal tissues, can cause such widespread biological effects, ranging from perturbing local mitochondrial metabolic mechanisms to affecting extramitochondrial signalling events, transcription and protein expression.

## Materials and Methods

### Chemicals and Solvents

Acetonitrile (ACN), Potassium chloride (KCl), Sodium chloride (NaCl), Potassium phosphate monobasic (KH_2_PO_4_), Deuterium oxide (^2^H_2_O), Disodium phosphate (Na_2_HPO_4_), _S_odium azide (NaN_3_) and 3-(trimethylsilyl) propionate-2,2,3,3-*d*_4_ sodium salt (TSP) were obtained from Sigma-Aldrich, United Kingdom. Deionized water was purified using an in-house Synergy Type 1 Ultrapure Water System from Milli-Q^R^ (Merck). All solvents were HPLC graded (Fisher Scientific).

### Animal models

A total of 30 male C57BL6/J laboratory mice, each weighing 25g were used for this study. Cyclophilin D-deficient mice (KO) were generated by targeted disruption of the *ppif* gene20 and provided by Dr Derek Yellon (University College London, UK) and Dr Michael A. Forte (Oregon Health and Sciences University, USA). Animal care and experimental protocols and procedures was approved by the University of Liverpool Biomedical Services Unit (BSU). All experiments comparing WT and *ppif*−/− (KO) mice were conducted using a total of 30 male C57BL6/J laboratory mice. Each cohort contained five biological replicates for all biological matrices. After 7 days of acclimatisation and feeding on rat chow diet and water, mice tissues (liver, heart, and pancreas) and blood were immediately collected after sacrifice. and placed on ice. Serum (0.8 mL) was then collected into 1.5 mL pre-washed (with phosphate buffer (PBS) pH 7.4) microfuge tubes (collection tubes that uses gel to separate blood cells from plasma was avoided because EDTA and other stabilizers give additional signals in the NMR spectra). Tubes containing serum were prepared and stored according to standard site-specific procedures. 50 mg of tissues were sectioned into small pieces using sterile scalpel and forceps, individually placed into 1.5 mL microfuge tubes and immediately flash frozen in liquid nitrogen to arrest any enzymatic or chemical reactions. Frozen samples were preserved at −80°C until metabolite extraction.

### Metabolite Extraction, Lysate Clarification, and Solvent Lyophilisation

Frozen tissues were suspended in 0.8mL of ice-cold 50:50 (v/v) acetonitrile/water containing 0.2mM TSP as an internal NMR reference. Samples were sonicated in an ice bath *via* three cycles of 30s pulse (with 30s rest) at amplitude 6.2μm using an exponential probe frequency of 23kHz. The sonicator probe was cleaned with ethanol followed by distilled water between each sample to avoid cross-contamination. All lysed homogenates were centrifuged at 16,000*g* for 10min at 4°C to separate the tissue particulates. Supernatants were transferred to fresh cryotubes and flash frozen by submerging in liquid nitrogen for a minimum of 10s. Frozen samples were lyophilised with an Heto PowerDry LL3000 freeze dryer at −55°C for 10 hours to remove the solvent. Samples for this study were stored at −80°C for no more than 7 days before use.

### Sample Preparation

All lyophilised samples were resuspended in 200μL of NMR sample buffer (100mM Na_2_HPO_4_ and 1.2mM NaN_3_ in ^2^H_2_O, pH 7.4). Samples were vortexed and then centrifuged at 21,500g for 15min at 20°C. 190μL of supernatant was transferred into 3mm outer diameter SampleJet NMR tube for data acquisition. Serum samples were thawed and 100 μL added to NMR sample buffer (100mM Na_2_HPO_4_ in ^2^H_2_O, pH 7.4 and 1.2mM NaN_3_ in H2O). Samples were first vortexed and then centrifuged at 13,000g for 2 mins at 20°C. 198 μL of supernatant was transferred into a 3 mm outer diameter SampleJet NMR tube for data acquisition.

#### NMR Data Acquisition

### Tissue extract

All ^1^H NMR spectra were acquired at 25°C (± 0.1ºC) using a Bruker Avance III 700 MHz spectrometer equipped with a [^1^H, ^15^N, ^13^C]-TCI Cryoprobe. The field frequency was locked on ^2^H_2_O. One-dimensional ^1^H experiments were acquired using a nuclear Overhauser effect spectroscopy (NOESY) presaturation sequence (RD−90°-*t*−90°-*t*_m_−90°-ACQ) to suppress the residual water peak and Carr-Purcell-Meiobom-Gill (CPMG) pulse sequence (RD−90°-(*t*−180°-*t*)_n_-ACQ) to observe low molecular weight metabolites during a total spin-spin relaxation delay of 4s. For optimal signal-to-noise, spectra were acquired with 256 scans and 48K points, with a spectral width of 30ppm. All spectra were processed identically using an automated routine in the (Bruker Topspin 3.5) software for line broadening, auto-phasing, (followed where necessary by manual phasing), baseline correction, and referencing to TSP (δ; chemical shift (0.0 ppm)

### Serum sample

^1^H NMR spectra of serum samples was acquired at 37°C (± 0.1ºC) using the 700 MHz spectrometer as described for the tissue extracts. The field frequency was locked on 90% H_2_O and 10% ^2^H_2_O. 1-D NOESY and 1-D CPMG experiments were applied to serum samples during a total spin-spin relaxation delay of 3s. Spectra were acquired with 256 scans and 73K points, with a spectral width of 17 ppm. All spectra were processed identically with 0.3 Hz line broadening, auto-phasing (followed where necessary by manual phasing), baseline-correction and referenced to glucose (δ 5.24) using an automated routine in the (Bruker Topspin 3.5) software.

### Statistical Analysis

Each ^1^H NMR spectrum was subdivided into a set number of 483 integrated regions called bins, that represent the relative number of protons resonating in the integrated region using Bruker Amix 3.9.14 software package. The region around the residual water resonance (4.5 - 4.8 ppm) was excluded and the total summation of signal intensities in each bin was calculated by relative intensities to reference region (TSP). Approximately 88 and 45 metabolites were respectively, annotated for tissue and serum samples by algorithmically auto-fitting spectral data sets using Chenomx profiler v 8.2 and verified using the Human Metabolome Database (https://hmdb.ca/), Biological Magnetic Resonance Data Bank (https://bmrb.io/), and in-house library 1D and 2D NMR spectra of selected compounds. Metaboanalyst 4.0 software package (http://www.metaboanalyst.ca/) was used to pre-process all data by normalising all samples to the concentration of the internal standard reference sample (0.2mM TSP) and scaling all features by auto-scaling (mean-centering and then dividing by the standard variation of each variable) to give each sample equal variance (i.e. to have an equal chance of contributing to the model). The concentration of each metabolite from individual tissues were superimposed into relative changes compared with the WT values, to allow one control comparison for all 3 tissues. Principal Component Analysis (PCA) was then carried out on normalised data to examine the metabolic profile to reflect the overall changes following the loss of CypD in mice serum, liver, heart, and pancreas. Statistical significance was determined by q values (false discovery rate corrected *P* values) for each metabolite (q < 0.05) is considered significant^38,39^. Meta-analysis was also carried out on the mice tissues using “Meta-analysis module” in Metaboanalyst 4.0, to identify consistent and robust metabolites that were altered in response to the loss of CypD in mice tissues. Individual dataset from the tissues were integrated by processing each data in consistent steps. Linear models (Limma) was performed on the normalised data, using the Fisher’s combined probability test at a p-value (FDR) cut off of 0.05 and fold change threshold of 1.5.

### Metabolomics Pathway Analysis

Pathway Analysis module on Metaboanalyst 4.0^40^, was used to evaluate the impact of metabolites that were consistently altered in a significant manner following the loss of CypD in mice tissues and the impact of significant differential metabolite from the serum samples on different metabolic pathways. Fisher’s exact test was used to perform the algorithm for the revelation of the metabolic pathways that were altered in response to deletion of CypD in mice serum and tissues, while the compound importance in the given metabolic pathway was estimated through its betweenness centrality and out of degree centrality (quantitative measure of the position of a node relative to the other nodes)^40^. The impact of the pathway was calculated as the sum of the importance measures of the matched metabolites normalized by the sum of the importance measures of all metabolites in each pathway^47^. Statistical significance for each impacted metabolic pathway in mice tissues and serum, was determined by q values (false discovery rate corrected *P* values) and q ≤ 0.05 was considered to be significant^38,39^. To further assess the impact of CypD deficiency on mice tissues and serum, pathway analysis using “Metabolomics pathway analysis” (MetPA) software package from Metaboanalyst 4.0^42^, was performed on metabolites that were consistently altered between CypD WT and KO cohorts, at a statistical significance value ≤ 0.05.

## Data Availability

The datasets generated and analysed during the current study are available from the corresponding author on reasonable request

## Acknowledgements

This study was supported by the postgraduate bursary from the University of Liverpool. The MRC is acknowledged for its funding of the 700 MHz spectrometer (MRC Grant No: MR/M009114/1).

## Author Contributions

Y.I.A carried out the research, data analysis and wrote the manuscript, O.S.A co-wrote the manuscript, supported with the experiments and data analysis, Y.O supported with the experiment, D.N.C and RS co-designed the study, L-YL designed the study and wrote the manuscript.

