## Supplementary Tables for "Untargeted ^1^H NMR Metabolomics and Pathway Analysis Reveals Dysregulated Proteostasis in Cyclophilin D (CypD)-deficient Mice Tissues"

### Contents:

1. Supplementary Table 1: List of identified and quantified metabolites in CypD deficient mice serum and tissue extract.
2. Supplementary Table 2: List of altered metabolic pathways in CypD deficient mice (a) tissues and (b) serum.
3. Supplementary Figure 1: Percentage of metabolite perturbation by CypD deficiency in mice tissues and serum.

### Supplementary Information

**Metabolite Identification and Quantification:** Metabolites extracted from each biological matrix were identified and quantified using the Chenomx Profiler module v 8.2. To identify the compounds in each sample, the appropriate proton signals arising from different mixture of metabolites in each sample were matched with the hundreds of reference compound signatures in Chenomx library. The software automatically adjusts the reference library, using a series of algorithm, to reflect the sample and acquisition conditions (e.g. pH, NMR field strength, spectra line width, and temperature). Peak intensity and location in each spectrum (sample) were then manually adjusted for fine-tuned concentration values. Individual metabolites were also further verified from various sources, including earlier published articles, literatures and cross checked from the Human Metabolome Database (HMDB), Biological Magnetic Resonance Bank (BMRB) and in-house compound 1D and 2D standards. Peak assignment was also validated with $^{1}$H-$^{13}$C Heteronuclear Single Quantum Coherence (HSQC) and $^{1}$H-$^{1}$H Correlation Spectroscopy (COSY). Fold changes and statistical significance value for each identified metabolite were determined by Independent Samples t.Test and Fold Change analysis in Metaboanalyst v 4.0. The list of identified metabolites along with their chemical shifts, mean fold changes and statistical significance of the Cyclophilin D Knockout (CypD KO) relative to the wild-type (CypD WT), are provided in Supplementary Table 1.

**Supplementary Table 1:** List of identified metabolites in CypD deficient mice tissue and serum extract using $^{1}$H NMR spectroscopy.

| No | Metabolite | δ $^{1}$H (ppm) | FC (LV) | FC (PC) | FC (HT) | FC (SE) |
|---|---|---|---|---|---|---|
| 1 | 2-oxoglutarate | 3.01 (t) | 1.77* | 1.43† | 1.58 † | ------ |
| 2 | 3-hydroxybutyrate | 1.22 (d) | 1.97* | 1.62† | 1.50‡ | 1.95‡ |
| 3 | 3-methylhistidine | 7.05 (s) | 2.11* | 2.27† | 0.81† | 1.53‡ |
| 4 | Acetate | 1.90 (s) | 0.58 | 1.61† | 0.98 | 0.93 |
| 5 | Acetoacetate | 2.27 (s) | 1.74* | 1.41† | 1.61* | 1.40‡ |
| 6 | Acetylcarnitine | 3.18 (s) | 0.47 | 0.40* | 0.35* | 0.77† |
| 7 | Adenine | 8.17 (s) | 1.75† | 2.27† | 1.73* | ------ |
| 8 | Adenosine | 8.33 (s) | 1.49 | 1.05 | 0.63‡ | ------ |
| 9 | ADP | 8.53 (s) | 1.30* | 1.04 | 1.41* | ------ |
| 10 | Agmatine | 3.01 (t) | 3.03* | 2.69† | 1.99† | ------ |
| 11 | Alanine | 1.47 (d) | 2.25* | 1.62† | 1.52† | 1.50* |
| 12 | AMP | 8.60 (s) | 2.03* | 3.63* | 1.74* | ------ |
| 13 | Arginine | 1.91 (m) | 2.85* | 1.66† | 1.74† | 1.59† |
| 14 | Asparagine | 2.85 (dd) | 1.91† | 2.00† | 1.85† | ------ |
| 15 | Aspartate | 2.84 (dd) | 1.72* | 1.65† | 2.82‡ | 1.87† |
| 16 | ATP | 8.50 (s) | 0.77* | 0.82‡ | 0.67‡ | ------ |
| 17 | β-alanine | 3.17 (t) | 1.65‡ | 2.35† | 1.69† | 3.02† |
| 18 | Betaine | 3.25 (s) | 1.50‡ | 1.60‡ | 2.85† | 0.80 |
| 19 | Carnitine | 3.22 (s) | 2.20† | 1.02 | 0.64† | ------ |

| | | | | | | |
|---|---|---|---|---|---|---|
| 20 | Choline | 3.19 (s) | 2.19‡ | 1.82 † | 1.66† | 0.79‡ |
| 21 | Citrate | 2.54 (d) | 0.73* | 0.45‡ | 1.29† | 0.85 |
| 22 | Creatine | 3.02 (s) | 1.45* | 0.68‡ | 0.69† | ------ |
| 23 | Creatinine | 3.03 (s) | ------ | ------ | ------ | 1.50‡ |
| 24 | Dimethylamine | 2.71 (s) | 0.54† | 0.44 | ------ | ------ |
| 25 | Dimethylglycine | 2.91 (s) | 3.04* | 0.55† | 1.11 | 0.79‡ |
| 26 | Ethanolamine | 3.13 (m) | 0.45† | 0.78† | 0.62 | ------ |
| 27 | Formate | 8.44 (s) | 0.27† | 1.16‡ | 0.66‡ | 0.62† |
| 28 | Fumarate | 6.50 (s) | 1.73‡ | 1.23 | 1.20† | 0.76 |
| 29 | Glucose | 5.22 (d) | 1.86‡ | 1.46‡ | 1.69‡ | 0.73‡ |
| 30 | Glucose-1-phosphate | 5.45 (dd) | 1.71* | 1.13 | 1.25 | ------ |
| 31 | Glucose-6-phosphate | 4.63 (dd) | 1.28‡ | 3.49* | 2.10* | ------ |
| 32 | Glutamate | 2.34 (m) | 2.66* | 2.19† | 2.50† | 0.74† |
| 33 | Glutamine | 2.44 (m) | 2.74* | 2.58‡ | 1.95† | 0.26† |
| 34 | Glutathione (reduced) | 2.57 (m) | 2.43* | 2.08‡ | 1.21† | ------ |
| 35 | Glutathione (oxidised) | 2.14 (m) | 0.78* | 0.84 | 1.03 | ------ |
| 36 | Glycerol | 3.55 (dd) | 2.44† | 1.82 † | 1.91† | 1.59† |
| 37 | Glycine | 3.56 (s) | 1.94† | 1.63‡ | 1.55‡ | 1.21 |
| 38 | GTP | 5.93 (d) | 1.40* | 1.80* | 1.93† | ------ |
| 39 | Guanosine | 7.99 (s) | 1.94† | 1.67‡ | 0.70† | ------ |
| 40 | Histamine | 7.95 (s) | 2.11* | 1.37† | 0.63† | ------ |
| 41 | Histidine | 7.05 (s) | 0.53† | 1.24† | 0.67† | 0.78* |
| 42 | Homocysteine | 3.87 (q) | 1.35* | 1.58‡ | 2.62† | ------ |
| 43 | Hypoxanthine | 8.18 (s) | 2.00* | 1.86† | 1.28‡ | ------ |
| 44 | IMP | 8.57 (s) | 1.80* | 1.94† | 0.74† | ------ |
| 45 | Inosine | 8.23 (s) | 1.82* | 3.40† | 2.11† | ------ |
| 46 | Isoleucine | 1.00 (s) | 1.97† | 1.71† | 1.85* | 0.60* |
| 47 | Lactate | 1.32 (d) | 0.37* | 0.74‡ | 0.54 † | 0.42* |
| 48 | Leucine | 0.95 (s) | 2.03* | 2.54† | 1.67* | 0.72* |
| 49 | Mobile lipid -CH3 | 0.85 | ------ | ------ | ------ | 1.62‡ |
| 50 | Mobile lipid -N(CH3)3 | 1.34 | ------ | ------ | ------ | 1.47‡ |
| 51 | Mobile unsaturated lipids | 5.44 | ------ | ------ | ------ | 1.51‡ |
| 52 | Lysine | 1.72 (m) | 0.40* | 1.79† | 1.53* | 0.80* |
| 53 | Malate | 2.66 (dd) | 1.46† | 1.38† | 1.45* | ------ |
| 54 | Methionine | 2.64 (t) | 2.12* | 1.61* | 1.88* | 0.47* |
| 55 | Myo-inositol | 4.05 (t) | 2.22* | 4.11† | 3.23† | 0.65* |
| 56 | NAD | 9.33 (s) | 0.61† | 1.73‡ | 1.59‡ | ------ |
| 57 | NADH | 8.48(s) | 1.91* | 2.29† | 2.30* | ------ |
| 58 | NADP | 8.42 (s) | 0.81 | 1.38† | 1.35‡ | ------ |
| 59 | NADPH | 8.46 (s) | 2.13* | 1.52† | 1.41† | ------ |
| 60 | Niacinamide | 7.58 (m) | 2.41* | 1.96‡ | 1.18‡ | ------ |
| 61 | Ornithine | 3.05 (t) | 1.34‡ | 1.49† | 1.70‡ | ------ |
| 62 | Oxaloacetate | 2.38 (s) | 1.49* | 1.62‡ | 1.55† | ------ |
| 63 | Pantothenate | 0.92 (s) | 0.36 † | 1.61‡ | 1.04 | ------ |
| 64 | Phenylalanine | 7.35 (m) | 0.38† | 2.49† | 1.17 | 0.99 |
| 65 | Phosphocholine | 3.21 (s) | 1.90† | 2.59† | 1.06 | 1.27‡ |

| | | | | | | |
|---|---|---|---|---|---|---|
| 66 | Phosphocreatine | 3.03 (s) | 0.32* | 1.52† | 1.64* | ------ |
| 67 | Phosphoenolpyruvate | 5.38 (t) | 1.52* | 2.21† | 2.19† | ------ |
| 68 | Phosphoethanolamine | 3.97 (m) | 1.60† | 1.74* | 0.65† | ------ |
| 69 | Proline | 3.32 (m) | 1.67* | 0.92 | 1.04 | 0.95 |
| 70 | Propionate | 1.04 (t) | 0.78 | 1.50† | 0.84 | ------ |
| 71 | Putrescine | 1.76 (m) | 1.38† | 2.49† | 1.27† | ------ |
| 72 | Pyruvate | 2.36 (s) | 1.86† | 2.45† | 1.70* | 1.38† |
| 73 | s-adenosylhomocysteine | 8.34 (s) | 2.10† | 2.03† | 2.14† | ------ |
| 74 | Sarcosine | 2.72 (s) | 0.36* | 1.10 | 1.06 | ------ |
| 75 | Serine | 3.95 (m) | 2.55* | 1.98† | 2.38* | 1.38* |
| 76 | sn-G3P | 3.22 (s) | 1.43† | 2.96† | 1.92† | ------ |
| 77 | Succinate | 2.39 (s) | 2.14* | 1.71† | 1.84† | 1.25 |
| 78 | Taurine | 3.24 (t) | 2.15* | 3.12† | 1.24† | ------ |
| 79 | Threonine | 4.26 (m) | 0.62† | 1.15‡ | 1.84‡ | 1.08 |
| 80 | Trimethylamine-N-oxide | 3.25 (s) | 0.42* | 2.36 † | 1.97* | ------ |
| 81 | Tryptophan | 7.73 (m) | 0.44* | 2.61† | 1.43* | 0.82 |
| 82 | Tyrosine | 7.19 (m) | 1.39† | 0.82‡ | 1.30* | 1.31‡ |
| 83 | UDP-galactose | 5.99 (dd) | 2.04† | 1.26‡ | 0.97 | ------ |
| 84 | UDP-glucose | 5.63 (dd) | 1.54* | 1.18 | 1.26 | ------ |
| 85 | UDP-glucuronate | 5.62 (dd) | 2.83† | 1.41‡ | 1.58† | ------ |
| 86 | UDP-Glc-NAc | 5.55 (dd) | 2.08† | 1.30* | 1.59* | ------ |
| 87 | UDP-Gal-NAc | 5.42 (dd) | 1.62* | 1.26 † | 1.89† | ------ |
| 88 | Uracil | 7.53 (d) | 0.46‡ | 2.62† | 1.96† | ------ |
| 89 | Urea | 5.75 (s) | ------ | ------ | ------ | 2.63† |
| 90 | Uridine | 5.90 (dd) | 3.47† | 2.39‡ | 0.90 | 0.99 |
| 91 | Valine | 1.03 (d) | 2.52† | 2.63† | 1.74‡ | 0.69 |
| 92 | Xanthine | 7.89 (s) | 1.62† | 1.57† | 1.34 | ------ |

**Table 1**. Table displays the chemical shift, fold changes and FDR adjusted p-values of the individual compounds that were detected and identified in CypD deficient mice tissues and serum. Column 2 contains identified and quantified metabolites in CypD WT and KO mice serum and tissues. Column 3 displays the chemical shifts ($\delta$ $^1$H (ppm) region and the multiplicity (in bracket) for each metabolite. Fold changes (FC) of each metabolite in the KO Liver (LV), KO pancreas (PC), KO heart (HT), and KO serum (SE) relatively to their corresponding WT groups are, respectively, shown in columns 4, 5, 6, and 7. Metabolites with FC values < 1 and > 1 are, respectively, differentially lower and higher in the KO relative to the WT group. Undetected metabolites in each biological matrix are represented with (----). Statistical significance for each metabolite is represented with a statistical sign: *$p \le 0.0001$; †$p \le 0.001$; ‡$p \le 0.05$ and FC values with no statistical sign mean they are not significantly altered between groups. Abbreviations: ADP, Adenosine diphosphate; AMP, Adenosine monophosphate; ATP, Adenosine triphosphate; GTP, Guanosine triphosphate, IMP, Inosine monophosphate; NADH, nicotinamide adenine dinucleotide (NAD)+ hydrogen (H); NAD, nicotinamide adenine dinucleotide; NADPH, nicotinamide adenine dinucleotide (NAD)+ hydrogen (H) phosphate; NADP, nicotinamide adenine dinucleotide phosphate; sn-G3P, sn-glycero-3-phosphocholine; UDP-Gal-NAC, UDP-N-acetylgalactosamine; UDP-Glc-NAC, UDP-N-acetylglucosamine. FDR adjusted p-value < 0.05.

**Supplementary Information**

**Metabolic Pathways Analysis** Pathway Analysis module on Metaboanalyst 4.0 was used to evaluate the impact of metabolites that were significantly altered following the loss of CypD in mice tissues and serum. The significance of each altered metabolic pathway was determined by Fisher's exact test, at a q value (false discovery rate corrected *P* values) threshold ≤ 0.05 and the impact of the pathway was calculated as the sum of the importance measures of the matched metabolites normalized by the sum of the importance measures of all metabolites in each pathway. The list of significantly altered metabolic pathways alongside their q values and impact scores of CypD deficient mice tissues and serum, relative to their WT group, are provided in Supplementary Table 2 and Table 3.

**Supplementary Table 2a:** List of altered metabolic pathways in CypD deficient mice tissues.

| No | Metabolic Pathways | Total | Hits | q value | Impact |
|---|---|---|---|---|---|
| 1 | Warburg Effect | 49 | 14 | 2.04E-05 | 0.38 |
| 2 | Citric Acid Cycle | 26 | 10 | 3.26E-05 | 0.41 |
| 3 | Purine metabolism | 63 | 15 | 3.26E-05 | 0.18 |
| 4 | Transfer of Acetyl Groups into Mitochondria | 18 | 8 | 5.33E-05 | 0.90 |
| 5 | Betaine metabolism | 18 | 4 | 5.33E-05 | 0.43 |
| 6 | Ammonia Recycling | 25 | 9 | 8.13E-05 | 0.20 |
| 7 | Methionine Metabolism | 39 | 11 | 8.13E-05 | 0.20 |
| 8 | Arginine and Proline Metabolism | 48 | 12 | 0.000106 | 0.34 |
| 9 | Malate-Aspartate Shuttle | 7 | 5 | 0.000133 | 0.71 |
| 10 | Urea Cycle | 23 | 8 | 0.000236 | 0.23 |
| 11 | Gluconeogenesis | 30 | 9 | 0.000245 | 0.22 |
| 12 | Aspartate Metabolism | 34 | 9 | 0.000629 | 0.8 |
| 13 | Glycine and Serine Metabolism | 50 | 11 | 0.000629 | 0.21 |
| 14 | Pyruvate Metabolism | 37 | 9 | 0.001122 | 0.39 |
| 15 | Glutamate Metabolism | 45 | 10 | 0.001122 | 0.24 |
| 16 | Alanine Metabolism | 14 | 5 | 0.005154 | 1 |
| 17 | *Tyrosine Metabolism* | *55* | *10* | *0.005362* | *0.09* |
| 18 | *Carnitine Synthesis* | *16* | *5* | *0.009108* | *0* |
| 19 | Pterine Biosynthesis | 18 | 5 | 0.01539 | 0.365 |
| 20 | Valine, Leucine, and Isoleucine Degradation | 51 | 7 | 0.01539 | 0.19 |
| 21 | Histidine Metabolism | 35 | 7 | 0.015691 | 0.36 |
| 22 | Glycolysis | 20 | 5 | 0.021975 | 0.13 |
| 23 | Mitochondrial Electron Transport Chain | 15 | 4 | 0.03586 | 0.29 |
| 24 | Androstenedioe Metabolism | 23 | 5 | 0.03586 | 0.25 |
| 25 | Glycerolipid Metabolism | 23 | 5 | 0.03586 | 0.16 |
| 26 | *Ethanol Degradation* | *15* | *4* | *0.03586* | *0.03* |
| 27 | Folate Metabolism | 24 | 5 | 0.039348 | 0.21 |
| 28 | *Butyrate Metabolism* | *16* | *4* | *0.040983* | *0.05* |
| 29 | Glucose-Alanine Cycle | 9 | 3 | 0.042423 | 0.56 |

| | | | | | |
|---|---|---|---|---|---|
| 30 | Phenylalanine and Tyrosine Metabolism | 25 | 5 | 0.042423 | 0.22 |

**Table 2a.** Table displays the altered metabolic pathways and their q (FDR adjusted p) values of commonly altered metabolites in CypD deficient mice tissues. Column 2 contains the metabolic pathways that were commonly altered in response to the loss of CypD in mice tissues. Column 3 displays the total compounds present in each pathway according to Small Molecule Pathway Database (SMPDB). Column 4 displays the number of metabolite (from our data) found in each pathway. Statistical significance value and Impact score are shown in column 5 and column 6, respectively. Metabolic pathways that were italicised indicates that they are not biologically relevant due to their given impact score which < 1.0. Statistical significance and biological relevance of each altered metabolic pathways are respectively, determined by q ≤ 0.05 and impact score ≥ 1.0.

**Supplementary Table 2b:** List of altered metabolic pathways in CypD deficient mice serum.

| No | Metabolic Pathways | Total | Hits | q value | Impact |
|---|---|---|---|---|---|
| 1 | Urea Cycle | 23 | 4 | 0.002928 | 0.18 |
| 2 | Ammonia Recycling | 25 | 4 | 0.004024 | 0.24 |
| 3 | Beta-Alanine Metabolism | 26 | 4 | 0.00465 | 0.38 |
| 4 | Aspartate metabolism | 34 | 3 | 0.064414 | 0.6 |
| 5 | Histidine Metabolism | 35 | 3 | 0.069165 | 0.37 |
| 6 | Phosphatidylcholine Biosynthesis | 18 | 2 | 0.086417 | 0.17 |
| 7 | Arginine and Proline Metabolism | 48 | 3 | 0.144 | 0.16 |
| 8 | Ketone Body Metabolism | 12 | 2 | 0.28794 | 0.34 |
| 9 | Alanine Metabolism | 14 | 2 | 0.32739 | 1 |

**Table 2b.** Table displays the altered metabolic pathways and their q (FDR adjusted p) values of commonly altered metabolites in CypD deficient mice serum. Column 2 contains the metabolic pathways that were commonly altered in response to the loss of CypD in mice serum. Column 3 displays the total compounds present in each pathway according to Small Molecule Pathway Database (SMPDB). Column 4 displays the number of metabolites from the dataset in this study found in each pathway. Statistical significance value and Impact score are, respectively, shown in columns 5 and 6. Statistical significance and biological relevance of each altered metabolic pathways are respectively, determined by q ≤ 0.05 and impact score ≥ 1.0.

**Supplementary Information**

**Supplementary Figure 1:** Percentage of significantly perturbed metabolites by CypD deficiency in mice tissues and serum.

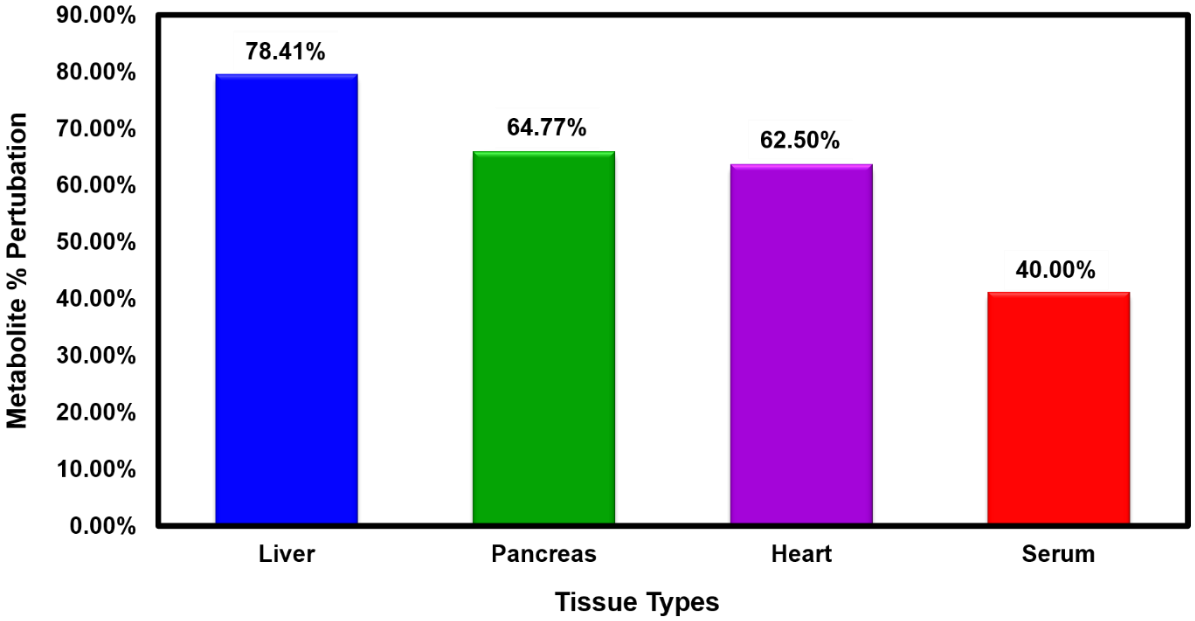


**Figure S1.** Percentage of significant metabolites perturbed by CypD deficiency in mice tissues and serum. This was calculated relative to the total metabolites detected in each tissue and serum samples. The liver demonstrated the highest impact of CypD deficiency by 78.41% while the serum was the least impacted by 40%.
